# Base-specific mutational intolerance near splice-sites clarifies role of non-essential splice nucleotides

**DOI:** 10.1101/129312

**Authors:** Sidi Zhang, Kaitlin E. Samocha, Manuel A. Rivas, Konrad J. Karczewski, Emma Daly, Ben Schmandt, Benjamin M. Neale, Daniel G. MacArthur, Mark J. Daly

## Abstract

Variation in RNA splicing (i.e., alternative splicing) plays an important role in many diseases. Variants near 5′ and 3′ splice sites often affect splicing, but the effects of these variants on splicing and disease have not been fully characterized beyond the 2 “essential” splice nucleotides flanking each exon. Here we provide quantitative measurements of tolerance to mutational disruptions by position and reference allele-alternative allele combination. We show that certain reference alleles are particularly sensitive to mutations, regardless of the alternative alleles into which they are mutated. Using public RNA-seq data, we demonstrate that individuals carrying such variants have significantly lower levels of the correctly spliced transcript compared to individuals without them, and confirm that these specific substitutions are highly enriched for known Mendelian mutations. Our results propose a more refined definition of the “splice region” and offer a new way to prioritize and provide functional interpretation of variants identified in diagnostic sequencing and association studies.

## Introduction

RNA splicing and alternative splicing are fundamental regulatory processes connecting transcription and translation. Splicing defects have been shown to make major contributions to the allelic architecture of numerous diseases including cystic fibrosis^1^, facioscapulohumeral muscular dystrophy^2^ and cancer^3^. Additionally, alternative splicing is particularly widespread in the nervous system and generates isoform diversity important to neuronal development and normal functioning^4^. For example, frequent misregulation of alternative splicing of microexons associated with splicing factor nSR100 is found in brains of patients with autism spectrum disorder^5^. A mouse model lacking nSR100 shows impaired neuronal development and inclusion of a nSR100-activated microexon was able to rescue part of the phenotype^6^.

Many previous studies have focused on how mutations affect splicing. Large-scale RNA-seq studies and smaller-scale minigene-based approaches have identified hundreds of eQTLs and sQTLs^7–10^, confirming a widespread influence of DNA variation on splicing variation. Based on these results as well as an understanding of cis- and trans-acting elements that affect splicing (such as branch site, polypyrimidine tract, and splicing enhancer and silencer motifs), computational algorithms have been developed to predict the effect of mutations on both general and tissue-specific splicing patterns^7, 11–13^. One important goal of studying mutations affecting splicing is to aid in the functional interpretation of the numerous single nucleotide variants (SNVs) identified in disease-mapping studies (both common alleles from GWAS and rare and *de novo* mutations found in Mendelian diseases and rare subtypes of common diseases). For instance, the GTEx Consortium demonstrated significant enrichment of sQTLs in ENCODE functional domains; Xiong et al. sifted through variants in disease candidate genes and prioritized mutations using the predicted likelihood that they will disrupt normal splicing, but did not find significant enrichment of splicing-disrupting variants in GWAS hits; and using a different method to identify sQTLs, Li et al. showed that the enrichment of sQTLs among GWAS SNPs is comparable or even larger in some cases than that of eQTLs^14^.

Of the variants confirmed or predicted to affect splicing, many are outside the two ultra-conserved (so-called “essential” or “canonical”) positions at both the 5′ (donor, typically GT) and 3′ (acceptor, typically AG) splice sites. Rivas et al. quantified the proportion of variants disrupting splicing based on RNA-seq results at +/-25bp from the splice sites^9^, clearly demonstrating signal beyond the essential sites.

The recent availability of the ExAC dataset^15^, a deep-coverage exome sequencing dataset with 60,706 individuals, permits a closer look at the near-splice site region since standard exome capture generally provides deep coverage 20-40 nucleotides to both sides of captured coding exons. Of particular importance, to assess the degree of mutational tolerance in different genes, Lek et al. developed an expectation-maximization approach to quantify the lack of protein-truncating variants compared to expectation in each gene (the probability of being loss-of-function intolerant, or pLI). As a result, genes with pLI value >= 0.9 (n = 3230, 17.7 percent) are particularly intolerant of disruptive mutations. Following this line of thought, we utilized the relative incidence of variation in splice regions of intolerant and tolerant genes to characterize the deleteriousness of mutations at each individual near-splice sites (see Material and Methods). Although different studies have focused on mutations in this region, none of them directly quantified the level of deleteriousness of mutations by both positions and reference/alternative allele types. Here we offer a systematic analysis of near-splice site mutations, identifying which sites are intolerant of mutation and confirming their impact through RNA-seq data and analysis of known Mendelian mutations, thereby developing a refined definition of “splice region” suitable for use in disease genetics.

## Material and Methods

We defined “near-splice site regions” as +/-10bp around the 5′ and 3′ splice sites. For all the following analysis, only canonical transcripts were considered (Gencode v19). See Figure S1 for additional nomenclature used throughout.

We used variation from the ExAC dataset (version 0.3, http://exac.broadinstitute.org) mapped to human reference genome (hg19) to scan for evidence of selection against variation in the near-splice site regions. First we tallied the number of A, T, C, and Gs in the reference genome at each near-splice site position across all canonical exons along with the number of variants observed in ExAC, correcting the numbers by coverage following previous methods^15^ in order to account for variants missed due to lower sequencing coverage. The corrected number of reference bases as well as number of mutations by reference and alternative allele are shown in Supplementary Table 1.

**Table 1.**
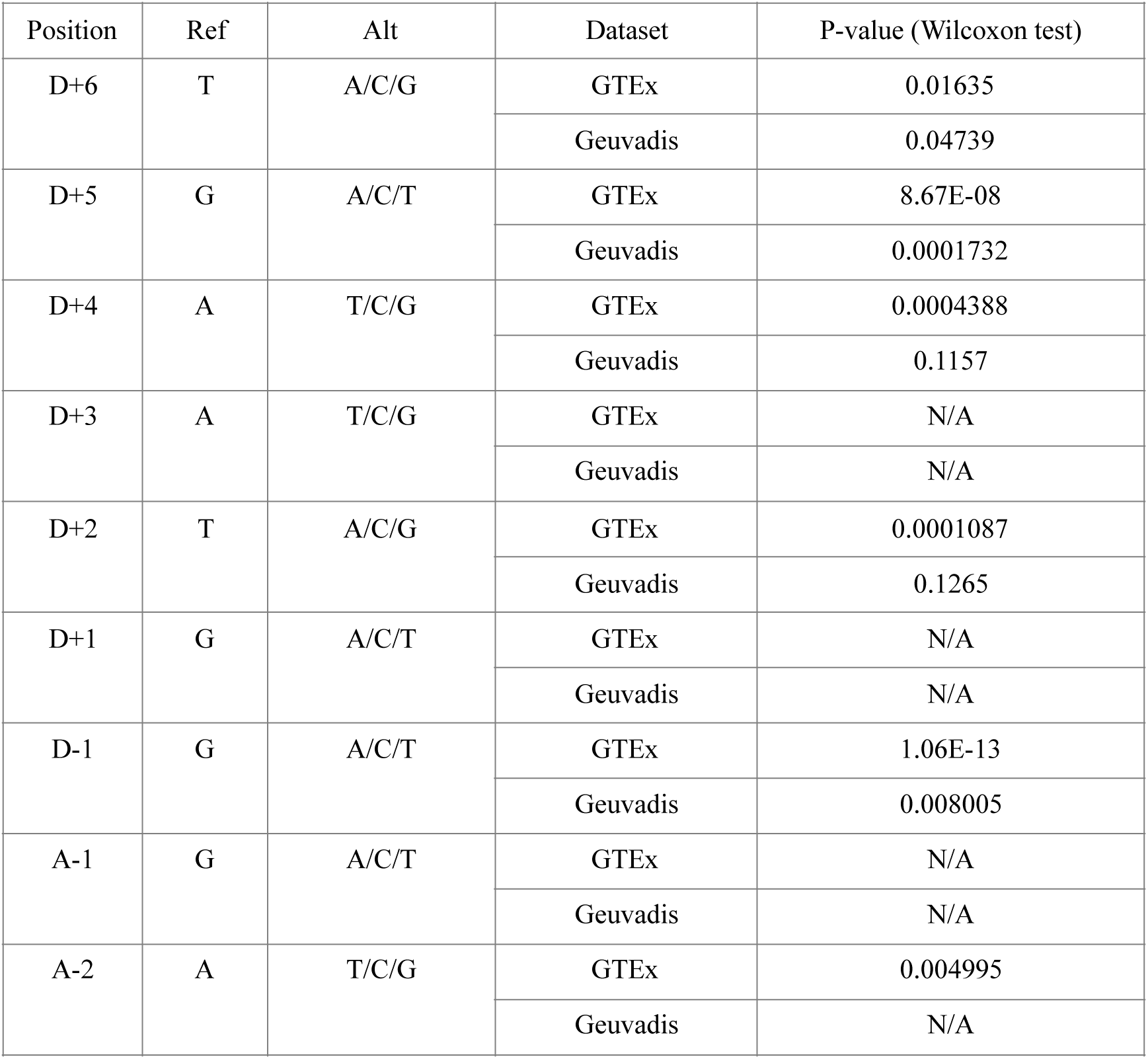
Comparison of the number of correct splicing reads between mutation carriers and homozygous reference individuals in GTEx and Geuvadis studies.

pLI, or the probability of a gene being loss-of-function intolerant, was developed in Lek et al. by comparing observed with expected rates of truncating mutations and identified that in 15-20% of genes, such mutations are under strong selection consistent with that seen in severe Mendelian haploinsufficiencies^15, 16^. Analyses that followed comparing loss of function intolerant (pLI >= 0.9, n = 3230) genes with others^15, 17, 18^ established that indeed heterozygous truncating mutations (nonsense, frameshift, and essential splice site mutations) in these genes often have significant medical consequences. We therefore theorized that any splice-region variants (beyond the essential splice sites) disrupting normal gene function should also be significantly depleted in loss-of-function intolerant genes. Thus we asked whether rates of mutations near splice junctions were significantly different between the same two groups of genes (LoF-tolerant and LoF-intolerant). We created a contingency table according to gene group (pLI < 0.9 versus pLI >= 0.9) and mutation count (number of bases with mutations versus number of bases lacking mutation, corrected for coverage) for each reference allele-alternative allele combination and calculated the Pearson’s Chi-squared statistic^19^, thus creating a 4×3 table of chi-squared statistics at each near-splice site position. We noticed that at some positions, the chi-squared statistics are consistently high for mutations from a specific reference allele regardless of the alternative allele to which they mutate; we also created a Pearson’s Chi-squared statistic for each reference base at all positions (Supplementary Table 1).

One caveat with comparing Pearson’s Chi-squared statistics of different reference allele-alternative allele combinations directly is that this statistic increases as counts in contingency tables increase (reflecting more power to detect distortions when sample size is larger). In other words, more common reference alleles tend to have higher statistics. Although the level of significance that we observe is well above what can be explained by commonness of alleles alone, we report odds ratios to better quantify the differences in mutation rates in the two groups of genes.

Since nucleotide context is such a major determinant of mutation rate, we employed the mutability-adjusted proportion of singletons (MAPS) as calculated in Lek et al. Briefly, the singleton ratio at each mutational site is adjusted by the mean singleton ratio of all mutations surrounded by the same 3-nucleotide sequence context. We compared MAPS of mutations changing reference alleles with particularly high Chi-squared statistics and odds ratios (named as constrained reference alleles) and other reference alleles in the splice region with that of ExAC missense and nonsense mutations, as well as ExAC missense mutations split by CADD categories, taken from Lek et al. directly.

RNA-seq analysis was carried out using GTEx and Geuvadis datasets. To test whether mutations identified as deleterious from mutational data alone have an effect on splicing, the median of normal^-^ ized splicing read counts (i.e., the proportion of RNA-seq reads spanning a splice junction that “correctly” connect adjacent exons) was compared between individuals carrying a mutation disrupting one of the constrained reference bases and individuals that are homozygous (for the constrained reference bases) by Wilcoxon rank sum test. A “N/A” result was reported when there was no enough data to perform the test.

The ClinVar dataset was downloaded from http://www.ncbi.nlm.nih.gov/clinvar/docs/maintenance_use/ in February 2016. Excesses of disease-causing variants were calculated using variants with clinical significance categories “Likely pathogenic”, “Pathogenic”, “Risk factor” and “Association”. We tested for the enrichment of mutations changing constrained reference alleles in ClinVar variants with a binomial test comparing proportions of mutations with constrained reference alleles with that in the general population (ExAC, including both pLI>=0.9 and pLI<0.9 genes). A one-tailed test was used because the alternative hypothesis is that the true proportions of such mutations in ClinVar are higher than those naturally occurred in the general population. We also tested for enrichment in “Benign/Likely benign” categories as a negative control.

## Results

We first sought to define which near-splice site positions are important for normal splicing. Consistent with implications from earlier RNA-seq studies^9^, the distribution of the chi-squared statistics over +/- 10 bp around splice junctions shows that, in addition to the four “essential splice” nucleotides, positions D+3, D+4, D+5, D+6, D-1 and A+1 positions are very significantly intolerant of mutations. Chi-squared statistics of exonic regions are, as expected, on average higher than intronic regions due to background genic mutation, but even considering this background, A+1 and D-1 (the initial and terminal coding nucleotides in each exon) are unusually constrained (Supplementary Table 1).

Interestingly, both chi-squared statistics and odds ratios for the four reference alleles at each near-splice site position demonstrate that a specific reference allele is more intolerant of mutations than others at the same positions. Figure 1A and 1B shows that reference base T at D+6, G at D+5, A at D+4 and D+3, G at D-1, and G at A+1 are significantly less tolerant of mutational alteration than are the other three reference bases at those same positions. This difference is clearly evident in the odds ratio of the variation rate between tolerant and intolerant genes (Figure 1C and 1D) indicating the statistical excess is not simply a function of sample size. In the following analyses, we name these reference bases that are particularly sensitive to mutations as “constrained reference bases”.

**Figure 1.**
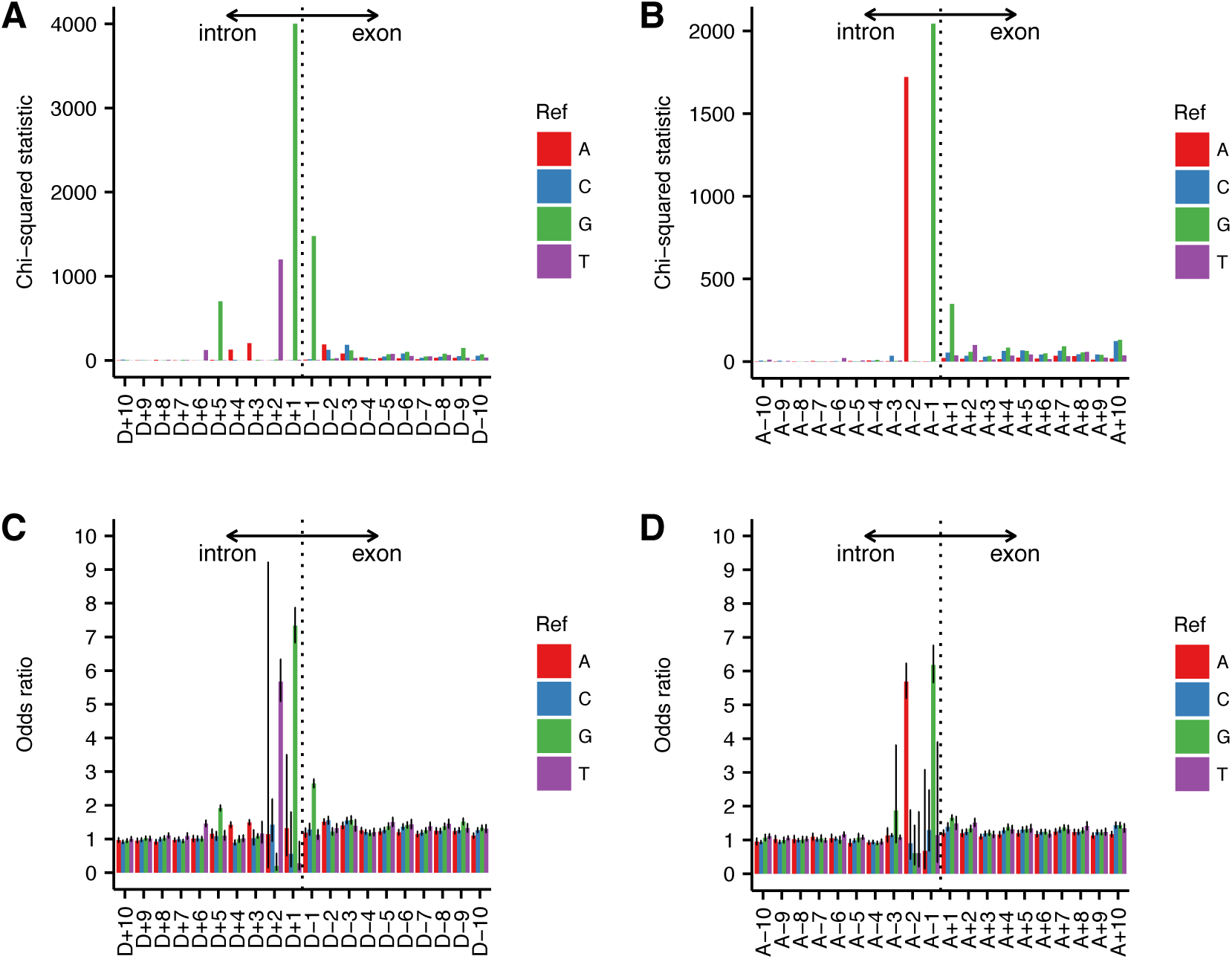
Quantification of intolerance to mutations split by positions and reference alleles. **A**, Chi-squared statistics at five prime end; **B**, Odds ratios at five prime end; **C**, Chi-squared statistics at three prime end; **D**, Odds ratios at three prime end.

Negative selection not only reduces the rate at which we find variants, but reduces the site frequency of observed sites when compared with neutral sites. To confirm the inference that selection is acting against specific nucleotide substitutions in the splice region, we first looked at the mutability-adjusted proportion of singletons (MAPS) as proposed in Lek et al. at constrained reference bases versus other three reference bases at the same positions (Figure 2). If mutations changing the constrained reference bases are indeed more deleterious, it is expected that there is a higher singleton ratio among these mutations because deleterious variants tend to be rarer. As we are comparing different reference bases, MAPS rather than the direct proportion of singletons is needed in order to account for systematic mutability differences given local nucleotide context. Consistent with the prediction from the chi-square analysis, MAPS is significantly higher at constrained reference bases than at other reference bases at the same positions. Compared to the average MAPS of functional classes reported in Lek et al., the constrained reference bases fall mostly between “missense variant” (0.0439) and “stop gained” (0.143), confirming their functional relevance. As a negative control, we also calculated MAPS for D+10 (a non-significant intronic site) as a comparator for the D+3 to D+6 sites and D-10 (an exonic site, which includes a mixture of missense and synonymous mutations) as a comparator for the D-1 and A+1 sites.

**Figure 2.**
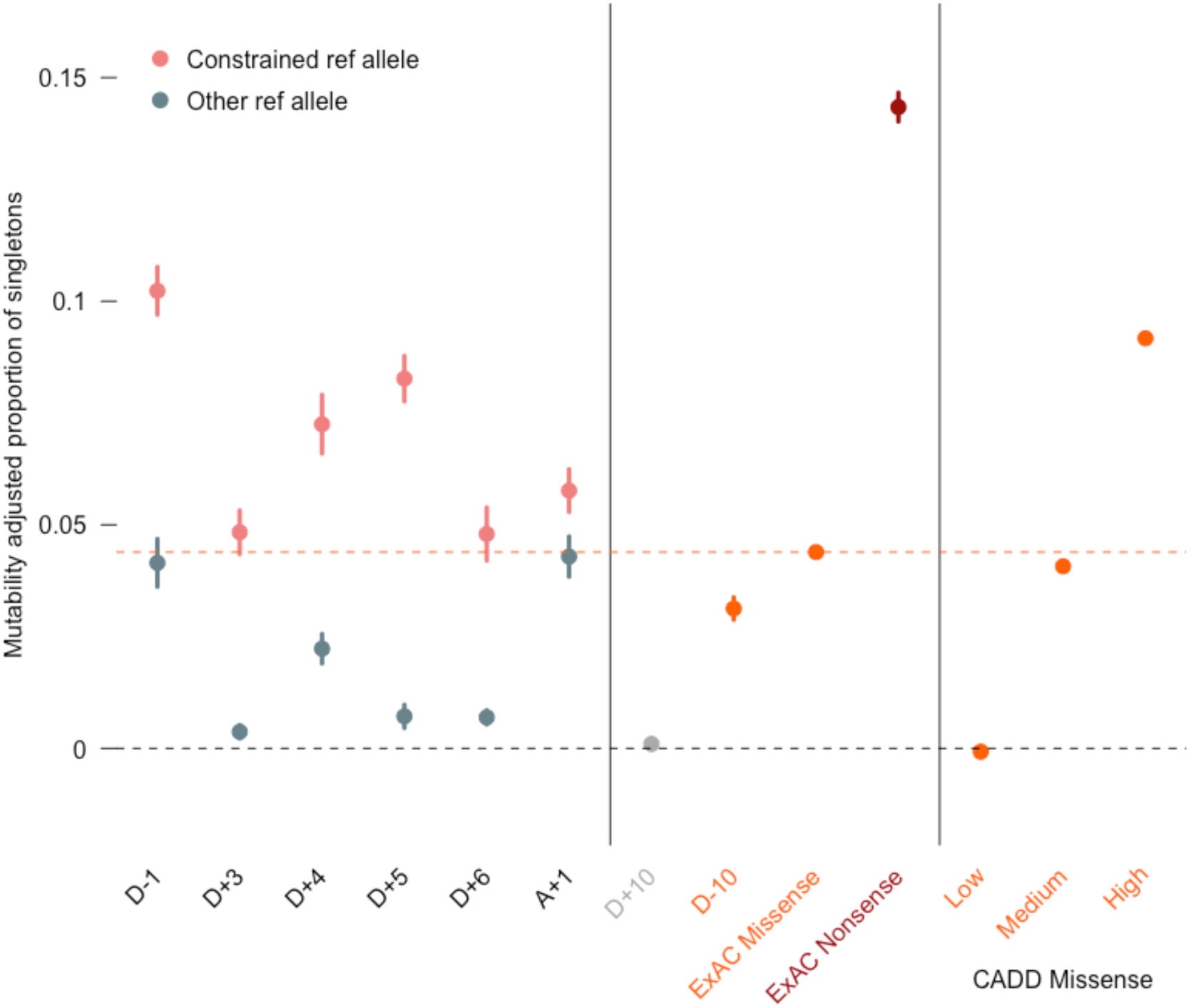
Singleton ratios adjusted by mutability at near-splice sites (leftmost panel) compared to ExAC missense, nonsense (middle panel) and missense split by CADD (rightmost panel) as references. “D+10” and “D-10” represent singleton ratios from a random intronic site and a random exonic site respectively and therefore serve as negative controls.

Given the clear evidence that natural selection does not tolerate substitutions in these “near-splice” regions in genes generally sensitive to heterozygous mutation, we then sought to characterize the functional effects on splicing and genetic impact on disease risk contributed by these variants. The availability of GTEx and Geuvadis data provides an excellent opportunity to examine the effect of mutations at near-splice sites on splicing^8, 20^. Specifically, we tested if heterozygous carriers of variants changing the constrained reference bases resulted in significantly less correct splice junctions reads compared to the homozygous reference allele carriers. Wilcoxon test shows that this is indeed the case for D+6 (Constrained ref: T; p = 0.016 in GTEx; p = 0.047 in Geuvadis); D+5 (Constrained ref: G; p = 8.67E-08 in GTEx; p = 0.0001732 in Geuvadis); D+4 (Constrained ref: A; p = 0.00044 in GTEx); D-1 (Constrained ref: G; p = 1.06E-13 in GTEx; p = 0.008 in Geuvadis) (Table 1). As a negative control, the same tests for unconstrained reference bases at the same locations do not show statistically significant result. These results strongly support the idea that mutations changing the constrained reference bases disrupt splicing and are less tolerated.

After confirming their impact on splicing, we next explored the impact these specific substitutions have on known, largely Mendelian, diseases. We found that mutations changing the constrained reference bases are significantly enriched in ClinVar variants with clinical significance categories “Likely pathogenic”, “Pathogenic”, “Risk factor”, or “Association”, but not in “Benign” or “Likely benign” categories (Table 2). “Likely pathogenic”, “Pathogenic”, “Risk factor”, and “Association” categories cover all curated variants with evidence of deleteriousness and together account for 39.3% of all ClinVar variants within 10bp of the splice sites. This result further support the idea that mutations changing the constrained reference alleles are more likely to be disease-related. Whenever possible, we also calculated enrichment p-values for other positions within the near-splice site region. There is a positive correlation between –log_10_(enrichment p-values) and chi-squared statistics/odds ratios from the above analysis, suggesting that the chi-squared statistics/odds ratios do indeed capture mutation deleteriousness as we think (Supplementary Table 2).

**Table 2.** Enrichment analysis of constrained reference base mutations in the ClinVar dataset.

| Position | Constrained ref mutation proportions in ExAC | Significance category “Pathogenic”, “Likely pathogenic”, “Risk factor” or “Association” |  |  | Significance category “Benign” or “Likely benign” |  |  |
| --- | --- | --- | --- | --- | --- | --- | --- |
|  |  | Constrained ref mutation count | Other ref mutation count | P-value (Binomial test) | Constrained ref mutation count | Other ref mutation count | P-value (Binomial test) |
| D+6 | 0.325 | 19 | 1 | 7.35E-09 | 19 | 75 | 0.997 |
| D+5 | 0.74 | 181 | 1 | <2.2E-16 | 21 | 22 | 0.999 |
| D+4 | 0.438 | 26 | 1 | 7.44E-09 | 5 | 44 | 1 |
| D+3 | 0.544 | 60 | 17 | 1.58E-05 | 24 | 18 | 0.42 |
| D-1 | 0.697 | 372 | 39 | <2.2E-16 | 83 | 170 | 1 |
| A+1 | 0.538 | 146 | 44 | 4.1E-11 | 70 | 140 | 1 |

These findings can be used to better annotate potential strong-acting mutations in established disease genes. As an example, we looked at a curated list of genes having more than three *de novo* protein-truncating variants each from the published studies of *de novo* variation in autism spectrum disorder, developmental delay (DD), and intellectual disability (ID), which have recently been jointly analyzed^18^. Thirteen genes also harbor at least one *de novo* near-splice site variants at the key positions identified above, indicating that these near-splice site variants may have a strong role in disease pathology (Supplementary Table 3). Additionally, six genes (*ZNF462*, *NCKAP1*, *NIN, EFTUD2, HNRNPK* and *KDM6B*) that have two protein-truncating variants each harbor a near-splice site variant at key positions, pushing these genes into the more convincing disease-related range.

In order to estimate the overall addition to disease mutation annotations by taking into account the constrained reference bases, we compared the number of credible deleterious mutations occurring at the essential splice sites versus the constrained reference bases at near-splice sites in ClinVar and the *de novo* variant datasets. Among ClinVar that meet the clinical significance category criteria mentioned above, 1284 and 973 mutations disrupt the 5′ and 3′ essential splice sites, respectively. By comparison, 806 mutations disrupt the six constrained references bases (D+3 A allele: 61; D+4 A allele: 27; D+5 G allele: 181; D+6 T allele: 19; D-1 G allele: 372; A+1 G allele: 146). In the *de novo* variant datasets including both autism and DD/ID individuals, 59 and 39 *de novo* variants occur at the 5′ and 3′ essential splice sites and a total of 35 mutations occur at the constrained reference bases in genes with pLI >= 0.9. Overall, about 36% more mutations in the splice region acquire new functional annotations if we take specifically intolerant splice junction mutations into account.

## Discussion

Variants outside splice sites are known to affect splicing, but a detailed evaluation of which mutations at non-essential sites near splice junctions affect splicing, and to what degree these contribute to disease, is still lacking. Our study provides a quantitative measure of the deleteriousness of mutations in near-splice sites, and shows that tolerance to mutations is reference allele specific. To validate this inference from the initial analysis, we showed that mutability-adjusted proportion of singletons (a site-frequency based metric correlated with negative selective pressure) at constrained reference bases are significantly higher compared to other reference bases at the same positions, and then further used RNA-sequencing data from Geuvadis and GTEx studies to show that mutations changing constrained reference bases resulted in significantly less correctly spliced reads. Importantly, we additionally confirmed that mutations changing the constrained reference bases are significantly enriched in ClinVar variants and make a meaningful addition to the allelic architecture of rare disease. In summary, by providing a detailed analysis of selective pressure and impact on splicing, we propose a refinement to the “splice region” definition suitable for use in Mendelian and complex disease exome analysis. As the specific pairings of positions and reference alleles have high impact on normal splicing if disrupted, they can therefore be used to prioritize and provide functional interpretations for mutations identified in association-type studies. By contrast, genome interpretation at present often annotates but quite frequently ignores the vastly larger, and largely benign, category of any variant in the ‘splice region’ within 10 or 20 bp of a splice junction – the annotation suggested here will enable a stronger consequence be attached to a much smaller number of variants. Specifically, surveying ExAC we find that an average participant sampled contains 11.17 variants in the splice region (within 10 bp of the splice junction) – only 0.87 of these are in the refined set of nucleotides and reference alleles listed here, and only 0.168 are in genes with pLI >= 0.9.

One limitation of our study is that we only considered variation at canonical transcripts. While we used canonical transcripts to capture overall deleteriousness of mutations at near-splice sites, to fully understand splicing in the context of specific diseases, it would be best to look at tissue-specific transcripts and isoforms. Additionally, the constrained reference alleles that we identified in this study are the most common reference alleles at the corresponding sites, and match the general “motif” at 5′ and 3′ splice sites proposed as early as 1987^21^ (Shapiro & Senapathy, 1987) based on commonness of alleles. However it is worth noting that our study uses a different set of methods and measures mutation deleteriousness rather than focusing on base composition frequencies alone.

## Supplemental Data

Supplementary data include one figure and one table in a PDF file and two separate excel tables.

## Acknowledgements

We would like to thank the ATGU community for their helpful discussions and insightful comments. KJK is supported by an NIGMS Fellowship (F32GM115208).

## References

1. Liu, X., Jiang, Q., Mansfield, S. G., Puttaraju, M., Zhang, Y., Zhou, W., Cohn, J. a, Garcia-Blanco, M. a, Mitchell, L. G., & Engelhardt, J. F. (2002). Partial correction of endogenous DeltaF508 CFTR in human cystic fibrosis airway epithelia by spliceosome-mediated RNA trans-splicing. Nature Biotechnology, 20(1), 47–52. https://doi.org/10.1038/nbt0102-47

2. Gabellini, D., D’Antona, G., Moggio, M., Prelle, A., Zecca, C., Adami, R., Angeletti, B., Ciscato, P., Pellegrino, M. A., Bottinelli, R., et al. (2006). Facioscapulohumeral muscular dystrophy in mice overexpressing FRG1. Nature, 439(7079), 973–977. https://doi.org/10.1038/nature04422

3. David, C. J., & Manley, J. L. (2010). Alternative pre-mRNA splicing regulation in cancer: Pathways and programs unhinged. Genes and Development, 24, 2343-2364. https://doi.org/10.1101/gad.1973010

4. Raj, B., & Blencowe, B. J. (2015). Alternative Splicing in the Mammalian Nervous System: Recent Insights into Mechanisms and Functional Roles. Neuron, 87, 14&27. https://doi.org/10.1016/j.neuron.2015.05.004

5. Irimia, M., Weatheritt, R. J., Ellis, J. D., Parikshak, N. N., Gonatopoulos-Pournatzis, T., Babor, M., Quesnel-Vallières, M., Tapial, J., Raj, B., O’Hanlon, D., et al. (2014). A highly conserved program of neuronal microexons is misregulated in autistic brains. Cell, 159, 1511–1523. https://doi.org/10.1016/j.cell.2014.11.035

6. Quesnel-vallières, M., Irimia, M., Cordes, S. P., & Blencowe, B. J. (2015). Essential roles for the splicing regulator nSR100 / SRRM4 during nervous system development. Genes and Development, 29, 746–759. https://doi.org/10.1101/gad.256115.114.4

7. Erkelenz, S., Theiss, S., Otte, M., Widera, M., Peter, J. O., & Schaal, H. (2014). Genomic HEXploring allows landscaping of novel potential splicing regulatory elements. Nucleic Acids Research, 42(16), 10681–10697. https://doi.org/10.1093/nar/gku736

8. GTEx Consortium. (2015). The Genotype-Tissue Expression (GTEx) pilot analysis: Multitissue gene regulation in humans. Science, 348(6235), 648–660. https://doi.org/10.1126/ science.1262110

9. Rivas, M. A., Pirinen, M., Conrad, D. F., Lek, M., Tsang, E. K., Karczewski, K. J., Maller, J. B., Kukurba, K. R., Deluca, D. S., Fromer, M., et al. (2015). Effect of predicted protein-truncating genetic variants on the human transcriptome. Science, 348(6235), 666–669.

10. Soukarieh, O., Gaildrat, P., Hamieh, M., Drouet, A., Baert-Desurmont, S., Frébourg, T., Tosi, M., & Martins, A. (2016). Exonic Splicing Mutations Are More Prevalent than Currently Estimated and Can Be Predicted by Using In Silico Tools. PLoS Genetics, 12(1), 1–26. https://doi.org/10.1371/journal.pgen.1005756

11. Barash, Y., Calarco, J. a, Gao, W., Pan, Q., Wang, X., Shai, O., Blencowe, B. J., & Frey, B. J. (2010). Deciphering the splicing code. Nature, 465(7294), 53–59. https://doi.org/10.1038/nature09000

12. Di Giacomo, D., Gaildrat, P., Abuli, A., Abdat, J., Frébourg, T., Tosi, M., & Martins, A. (2013). Functional analysis of a large set of brca2 exon 7 variants highlights the predictive value of hexamer scores in detecting alterations of exonic splicing regulatory elements. Human Mutation, 34(11), 1547–1557. https://doi.org/10.1002/humu.22428

13. Xiong, H. Y., Alipanahi, B., Lee, L. J., Bretschneider, H., Merico, D., Yuen, R. K. C., Hua, Y., Gueroussov, S., Najafabadi, H. S., Hughes, T. R., et al. (2015). The human splicing code reveals new insights into the genetic determinants of disease. Science, 347(6218). https://doi.org/10.1126/science.1254806

14. Li, Y. I., Van De Geijn, B., Raj, A., Knowles, D. A., Petti, A. A., Golan, D., Gilad, Y., & Pritchard, J. K. (2016). RNA splicing is a primary link between genetic variation and disease. Science, 352(6285), 600–604.

15. Lek, M., Karczewski, K. J., Minikel, E. V, Samocha, K. E., Banks, E., Fennell, T., O’Donnell-Luria, A. H., Ware, J. S., Hill, A. J., Cummings, B. B., et al. (2016). Analysis of protein-coding genetic variation in 60,706 humans. Nature, 536(7616), 285–291. https://doi.org/10.1038/nature19057

16. Cassa, C. A., Weghorn D., Balick D. J., Jordan D. M., Nusinow D., Samocha K. E., O’Donnell-Luria A., MacArthur D. G., Daly M. J., Beier D. R., et al. (2017). Estimating the selective effects of heterozygous protein-truncating variants from human exam data. Nature Genetics. https://doi.org/10.1038/ng.3831

17. Genovese, G., Fromer, M., Stahl, E. A., Ruderfer, D. M., Chambert, K., Landén, M., Moran, J. L., Purcell, S. M., Sklar, P., Sullivan, P. F., et al. (2016). Increased burden of ultra-rare protein-altering variants among 4,877 individuals with schizophrenia. Nature Neuroscience, 19(11), 1433–1441. https://doi.org/10.1038/nn.4402

18. Kosmicki, J., Samocha, K., Howrigan, D., Sanders, S., Slowikowski, K., Lek, M., Karczewski, K., Cutler, D., Devlin, B., Roeder, K., et al. (2017). Refining the role of de novo protein truncating variants in neurodevelopmental disorders using population reference samples. Nature Genetics, 49, 504–510. https://doi.org/doi:10.1038/ng.3789

19. Agresti, A. (2007). An Introduction to Categorical Data Analysis. Statistics. https://doi.org/10.1002/0471249688

20. Lappalainen, T., Sammeth, M., Friedländer, M. R., ’t Hoen, P. a C., Monlong, J., Rivas, M. a, Gonzàlez-Porta, M., Kurbatova, N., Griebel, T., Ferreira, P. G., et al. (2013). Transcriptome and genome sequencing uncovers functional variation in humans. Nature, 501(7468), 506–11. https://doi.org/10.1038/nature12531

21. Shapiro, M. B., & Senapathy, P. (1987). RNA splice junctions of different classes of eukaryotes: Sequence statistics and functional implications in gene expression. Nucleic Acids Research, 15(17), 7155–7174. https://doi.org/10.1093/nar/15.17.7155

